# AIFS: A novel perspective, Artificial Intelligence infused wrapper based Feature Selection Algorithm on High Dimensional data analysis

**DOI:** 10.1101/2022.07.21.501053

**Authors:** Rahi Jain, Wei Xu

## Abstract

**Background:** Feature selection is important in high dimensional data analysis. The wrapper approach is one of the ways to perform feature selection, but it is computationally intensive as it builds and evaluates models of multiple subsets of features. The existing wrapper approaches primarily focus on shortening the path to find an optimal feature set. However, these approaches underutilize the capability of feature subset models, which impacts feature selection and its predictive performance.

**Method and Results:** This study proposes a novel Artificial Intelligence infused wrapper based Feature Selection (AIFS), a new feature selection method that integrates artificial intelligence with wrapper based feature selection. The approach creates a Performance Prediction Model (PPM) using artificial intelligence (AI) which predicts the performance of any feature set and allows wrapper based methods to predict and evaluate the feature subset model performance without building actual model. The algorithm can make wrapper based method more relevant for high-dimensional data and is flexible to be applicable in any wrapper based method. We evaluate the performance of this algorithm using simulated studies and real research studies. AIFS shows better or at par feature selection and model prediction performance than standard penalized feature selection algorithms like LASSO and sparse partial least squares.

**Conclusion:** AIFS approach provides an alternative method to the existing approaches for feature selection. The current study focuses on AIFS application in continuous cross-sectional data. However, it could be applied to other datasets like longitudinal, categorical and time-to-event biological data.

## Background

Large feature space (*p*) is an important aspect of high dimensional data owing to the risk of model overfitting and poor model generalizability [1] and increased computational complexity [2, 3]. Feature selection is a solution which reduces the input feature space to smaller feature space (*q*) in a given dataset of sample size (*n*), which provides a parsimonious best fit model for the outcome, *y*.

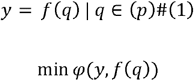

where, *f* represents the model function, and *φ* represents the error function. The approaches adopted for feature selection can be categorized into two groups. The first and simpler approach uses expert opinion for feature selection where features are selected using domain knowledge [4, 5] and allows feature selection before evaluating the data. This approach has limitation or no applicability if a feature has no or little availability of domain information, high dimensional feature space and/or presence of interactions among the features [6].

The second and prominent approach uses the sampled data to perform the feature selection which is broadly classified into filter, embedded and wrapper methods [7–9]. These methods could be used in supervised, semi-supervised or unsupervised learning frameworks [9–11]. Filter methods rely on the internal data structure of the features for selecting features. Commonly, information gain based techniques are used for univariate filtering of features [9, 12] and correlation based techniques are used for multivariate filtering of features [13]. They are computationally efficient, but interactions between the features may hinder the model performance. Embedded methods incorporate feature selection within the model building step by adding a penalization step in the model building process. They are efficient and have the ability to handle interactions between the features. LASSO based techniques [14–16] are commonly used for linear combination models, while tree-based algorithm [17] are used in non-linear combination models. Wrapper methods use an iterative approach where a model is built using a subset of features in which the performance is evaluated [18, 19]. The process is repeated until the best performance is obtained. It provides better performance than other methods, but it has a higher computational cost.

Most techniques have focused on reducing the computational cost of wrapper based methods by designing algorithms that reduce the optimization route to the target feature set *q*, i.e., using the minimum number of iterations to get *q*. The studies achieve this objective by focusing on the sampling of feature subset. Feature subset sampling step is commonly performed using either random sampling, sequential sampling or evolutionary sampling [20–23]. The random sampling approach arbitrarily generates the feature subset [20]. The sequential sampling approach adds or removes a feature sequentially from a feature set like forward sampling and backward sampling [18, 21]. The evolutionary sampling approach selects the feature subset based on the performance of features in the previous subset like genetic algorithm [22] and swarm optimization [23]. The number of iterations is an important bottleneck in improving the computation efficiency of the wrapper methods.

The wrapper methods assume that feature subset with target features should provide better performance than other feature subsets. Thus, the wrapper methods build models to estimate the performance for evaluation. The need to build a model for every single feature subset obtained in the sampling step creates another critical bottleneck in reducing computational complexity. Our research suggests that model building may not be the only approach to obtain performance value.

Currently, the existing wrapper methods partially or entirely discard the unselected models of feature subset in selecting the next population of feature subsets. Individually, each model may only be useful in providing performance information, but in combination, these models could help in identifying hidden relationships that could help in predicting the performance of unknown feature subset models. This may eliminate the need for building models for every single feature subset obtained in the sampling step. Accordingly, this study focuses on reducing the number of models that need to be built for a given number of feature subsets obtained in the sampling step of wrapper based feature selection.

In this study, we propose a novel Artificial Intelligence infused wrapper based Feature Selection (AIFS) algorithm. This algorithm can predict the performance of a feature subset using an existing artificial intelligence (AI) model rather than estimates the performance of a feature subset by building an actual AI model (like LASSO, Random Forest). AIFS is unique in many ways. Firstly, it is unique in its perspective as, unlike classical wrapper approaches of building models for every feature subset provided by feature subset sampling step, it builds models for only a fraction of the feature subset. Secondly, it provides a unique application of AI models, that are used to replace the AI model-based performance estimation step with AI model-based performance prediction step, which may reduce the computation time. Thirdly, AIFS is versatile, which allows its integration with existing statistical and machine learning techniques.

This paper provides the “Conceptual Framework” section to explain the basic framework of AIFS. The “Methodology” section explains the AIFS algorithm used in this paper. The algorithm performance is evaluated and compared against the existing feature selection methodologies for simulations and real studies in the “Simulation Studies” and “Real Studies” sections. Finally, we summarize and provide future directions for research in the “Conclusion and Discussion” section.

## Results

The performance of AIFS is evaluated and compared with standard methods like LASSO, adaptive LASSO, group LASSO, sparse partial least squares, elastic net and adaptive elastic net for both the simulated datasets and real data studies.

### Simulation Studies

We perform simulation studies to evaluate the proposed method and compare its performance with other feature selection methods. The study uses multivariate normal distributions to generate high-dimensional datasets for marginal and interaction models. The regression model, 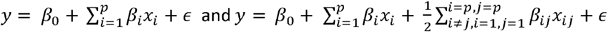 provides the outcome variable of the simulated data for marginal and interaction models, respectively. *ε ∼ N* (0, σ^2^), *x*_*i*_ *∼ N* (0,1) and {*x*_*ij*_} represents the pairwise interactions between features {(*x*_1_, *x*_2_), (*x*_1_, *x*_2_),,…, (*x*_p-1,_ *x*_p_)}. In the current study, only two-way interactions are considered for demonstration purposes, but it could be easily extended to higher-order interactions. Correlation is added between the first 15 features out of *p* marginal features using the covariance matrix as given below.

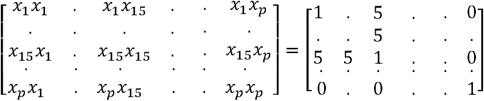

Multiple scenarios are created with the different number of noise features (Table 1). Non-zero value is assigned only to the true features. The AIFS approach is implemented both with and without a performance-based filter step. The final predictive model from selected features is prepared using either RIDGE regression (AIFS-LR) or non-penalized linear regression (AIFS-LLr). When no performance-based filter step is performed, model obtained from embedded feature selection stage is used as the final predictive model and is referred to as AIFS-L technique.

**Table 1:** Description of the simulation data

| Models | Scenario | $\beta$ (Non-Zero coefficients) | $p$ | Sample Size ($n$) | | $\sigma$ |
| --- | --- | --- | --- | --- | --- | --- |
|  |  |  |  | Train | Test |  |
| Marginal | 1_M | $\{\beta_i \mid i = \{1, \dots, 10\}\} =$<br>$\{0.5, -0.5, 0.5, -0.5, \dots, 0.5\}$ | 50 | 50 | 500 | 0.25 |
|  | 2_M |  | 50 | 100 | 500 | 0.25 |
|  | 3_M |  | 100 | 75 | 500 | 0.25 |
|  | 4_M |  | 100 | 100 | 500 | 0.25 |
| Interactions | 1_I | $\{\beta_i, \beta_{ij} \mid i = \{1, \dots, 10\}, j = i + 1, j < 11\} =$<br>$\{0.5, -0.5, 0.5, -0.5, \dots, 0.5\}$ | 15 | 100 | 500 | 0.25 |
|  | 2_I |  | 25 | 100 | 500 | 0.25 |
|  | 3_I |  | 50 | 100 | 500 | 0.25 |

### Computation Time estimation

We estimate computation time of the AIFS algorithm under different scenarios on a system with processor Intel® Core (TM) i7-8750H CPU@2.20GHz with 16 GB RAM on a Windows 10 64-bit operating system. The computation time is compared with the standard wrapper based approach that did not have the Performance Prediction Model (PPM). Since, standard wrapper (StW) does not have performance-based feature selection step, we compare it with AIFS-L method. Further, we add embedded feature selection step in StW. Thus, any performance difference is only associated with PPM model. Genetic algorithm is used to generate samples in feature subset sampling step with maximum number of iterations fixed to 200. Multiple scenarios are created for the comparative analysis of two algorithms (Table 1). The training datasets vary from 50-100 samples, while the test datasets contain 500 samples. In each scenario, training samples and test samples are independent samples that came from same distribution. Along with computation time, we evaluated both methods on their ability to select the target features and predictive performance of selected features. F1 score is used to determine the accuracy of selecting target features. Root Mean Square Error (RMSE) from the test data is used to determine the predictive performance of the model obtained from the embedded feature selection step. All the analysis is conducted using R 4.0.3 [24].

In both the marginal and interaction models (Table 2), AIFS consumed more time as compared to standard wrapper approach. This is counter intuitive, but this behavior is possible due to the PPM model upgradation step in AIFS. During each upgrade, sample size used for training PPM model increases. The current approach uses random forest to update PPM model and uses LASSO to build the base model. LASSO needs to build the model on a sample size of 50 or 100 but random forest needs to build a PPM model using at least 225 samples (Model 1_I) with sample size increasing during the execution of genetic algorithm.

**Table 2:** Wrapper methods comparison of computation time, target feature selection and predictive

| *Model* | *p* | *n* | **Performance** |  |  |  |  |  |
| --- | --- | --- | --- | --- | --- | --- | --- | --- |
|  |  |  | **Computation Time (minutes)** |  | **Target Feature Selection (F1 Score)** |  | **Predictive Performance (RMSE)** |  |
|  |  |  | *StW* | *AIFS-L* | *StW* | *AIFS-L* | *StW* | *AIFS-L* |
| 1_M | 50 | 50 | **7.57** | 24.85 | **0.48** | 0.47 | 0.55 | **0.43** |
| 2_M | 50 | 100 | **10.68** | 23.93 | **0.71** | 0.63 | 0.29 | **0.29** |
| 3_M | 100 | 75 | **11.30** | 18.22 | 0.29 | **0.33** | 0.64 | **0.48** |
| 4_M | 100 | 100 | **31.52** | 33.07 | 0.42 | **0.43** | 0.36 | **0.36** |
| 1_I | 15 | 50 | **0.97** | 2.18 | 0.41 | **0.73** | 1.20 | **0.38** |
| 2_I | 25 | 50 | **2.72** | 7.23 | 0.26 | **0.39** | 1.32 | **0.49** |

However, AIFS has a better or at par ability to discriminate between the target and noise features, especially for interaction models as compared to standard wrapper method. Similarly, predictive performance of the features shortlisted from AIFS is better or at par with standard wrapper method, especially for high dimensional data and interaction models. AIFS performance suggests that this methodology framework can be used as an alternative to the standard wrapper framework.

### AIFS comparison with standard methods

AIFS performance is compared with existing standard penalized regression methods namely LASSO, adaptive LASSO (ALASSO), group LASSO (GLASSO), elastic net (Enet), adaptive elastic net (AEnet) and sparse partial least squares (SPLS) in ten different trials. GLASSO is used only for interaction models. All the analysis is conducted using R 4.0.3 [24]. The standard methods are run using the inbuilt packages in statistical language R. *glmnet* package [25] is used for most methods except GLASSO and SPLS for which *glinternet* [26] and *spls* [27] packages were used. In the case of adaptive models, adaptive weights are obtained from ridge regression [28]. In the case of interaction models, all possible two-way interaction terms were created and entered the model. AIFS is implemented using the algorithm programmed in R.

The AIFS and the standard methods are evaluated on target feature selection and prediction performance. We evaluate the method’s ability to discriminate between true and noise features by measuring the selection of true features and rejection of noise features. We use RMSE from the test data as the predictive performance metric.

Table 3 shows the feature selection performance of different methods for marginal models. All methods have selected the targeted ten features which means that they can identify the target features in the marginal dataset. However, in most cases, the number of selected features is much higher, indicating that methods also select noise features. Compared to standard methods, the AIFS method selected a similar or lesser number of noise features which suggests that it has better discrimination ability between noise and target features than standard methods. Further, results from Figure 1 indicates better discrimination ability of the AIFS method than the standard methods. It is shown that frequency of selecting a noise feature is consistently lesser than the target features in all methods, but the maximum separation is found only for AIFS method. In addition, the area under curve (AUC) of the features was higher for AIFS method as compared to standard methods. Thus, in the case of marginal datasets, while all methods can identify the target features, AIFS outperforms all other methods with a lesser selection of noise features.

**Table 3:** Feature selection performance of different approaches in simulated scenarios for marginal models

| Scenario | Performance<br>(Number of<br>Features<br>Selected) | Target Features | Existing Models |  |  |  |  | AIFS |  |  |
| --- | --- | --- | --- | --- | --- | --- | --- | --- | --- | --- |
|  |  |  | *ALASSO* | *LASSO* | *SPLS* | Enet | AEnet | *AIFS-L* | *AIFS-LLr* | *AIFS-LR* |
|  |  |  | *Mean (Range)* |  |  |  |  |  |  |  |
| 1_M | Marginal<br>(p=50) | 10 | 24<br>(18-32) | 25<br>(18-37) | 23<br>(14-35) | 27<br>(18-36) | 26<br>(21-30) | 29<br>(24-33) | 15<br>(11-22) | **12**<br>**(10-16)** |
| 2_M | Marginal<br>(p=50) | 10 | 16<br>(11-35) | 23<br>(14-40) | 16<br>(10-39) | 25<br>(14-41) | 18<br>(11-35) | 24<br>(19-31) | 16<br>(10-31) | **12**<br>**(10-16)** |
| 3_M | Marginal<br>(p=100) | 10 | 27<br>(20-39) | 32<br>(16-57) | 25<br>(12-50) | 32<br>(21-45) | 28<br>(20-43) | 44<br>(29-59) | 18<br>(10-26) | **14**<br>**(10-21)** |
| 4_M | Marginal<br>(p=100) | 10 | 28<br>(14-46) | 33<br>(14-55) | 19<br>(11-47) | 32<br>(17-55) | 30<br>(15-48) | 44<br>(34-51) | 19<br>(10-45) | **13**<br>**(10-22)** |

**Figure 1.**
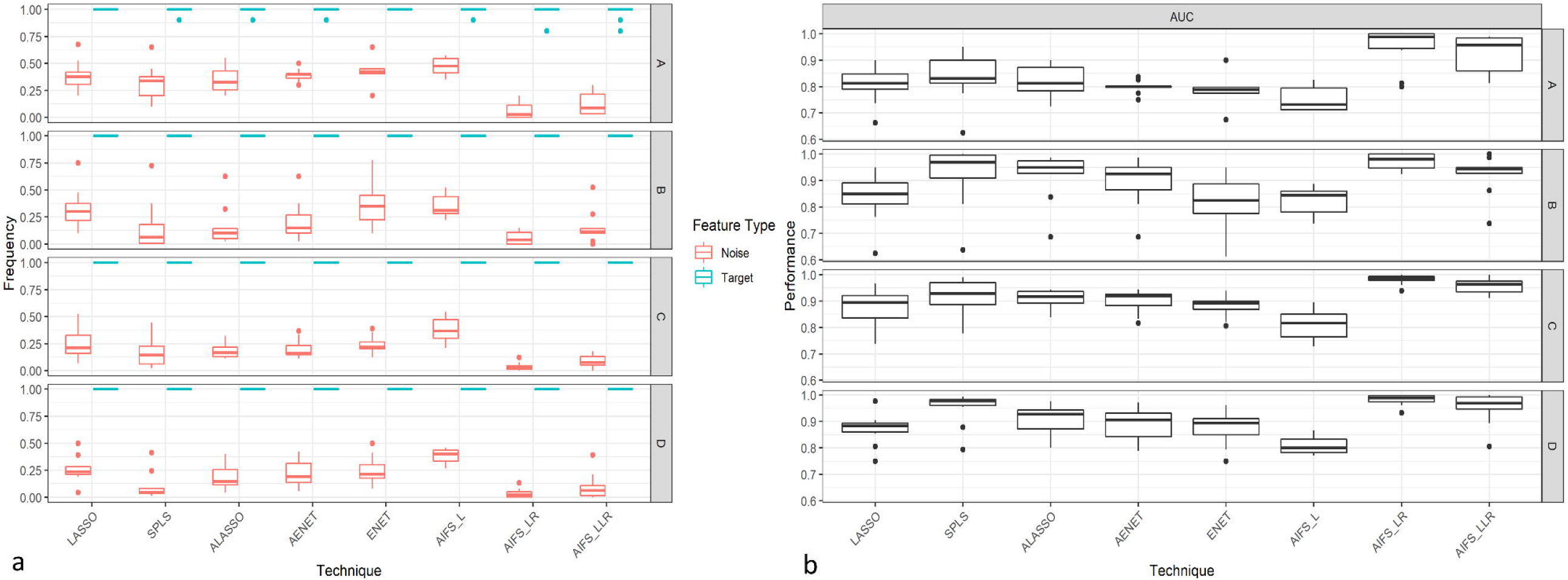
Comparison of different methods’ feature selection performance in marginal models a) Frequency of selection of target and noise features. b) AUC for predicting the target and noise features.

The results from the interaction models reiterate the results of the marginal scenario that the feature selection performance of AIFS is better or at par with the standard methods. Table 4 shows that like marginal models the number of features selected by all methods is more than the number of target features in most cases. This suggests that noise features are selected by all methods, but the number of noise features selected differs with methods. AIFS method selects a similar or lesser number of noise features compared to the standard methods, and results from Figure 2 suggest that AIFS may be selecting a lesser number of noise features compared to other methods. The results show that in low dimensional space, all methods can discriminate between the target and noise features by selecting the target features at a higher frequency as compared to noise features. However, in very high dimensions, only AIFS and GLASSO can perform. AUC performance of different methods also shows better or at par performance of AIFS as it can predict the target and noise features with greater or similar accuracy than other methods.

**Table 4:**
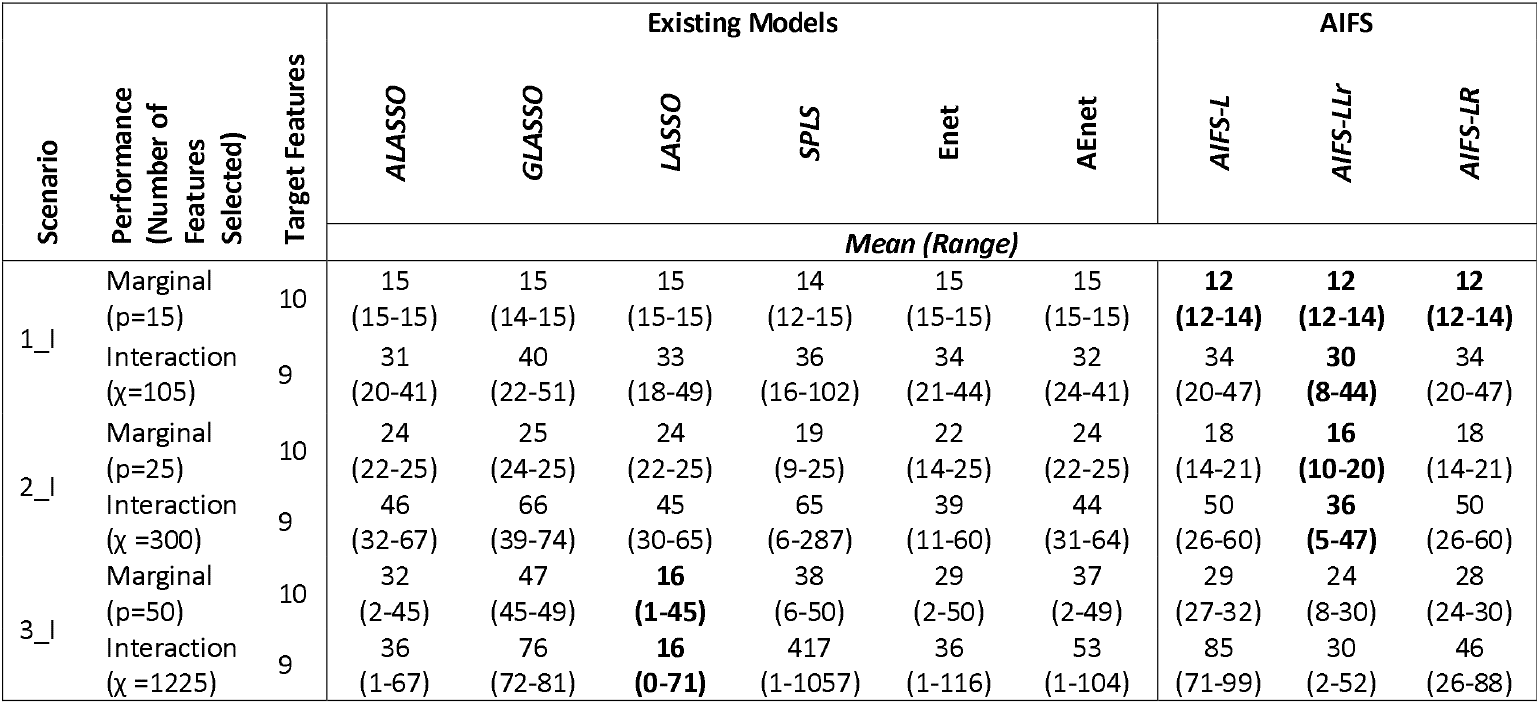
Feature selection performance of different approaches in simulated scenarios for interaction models

**Figure 2.**
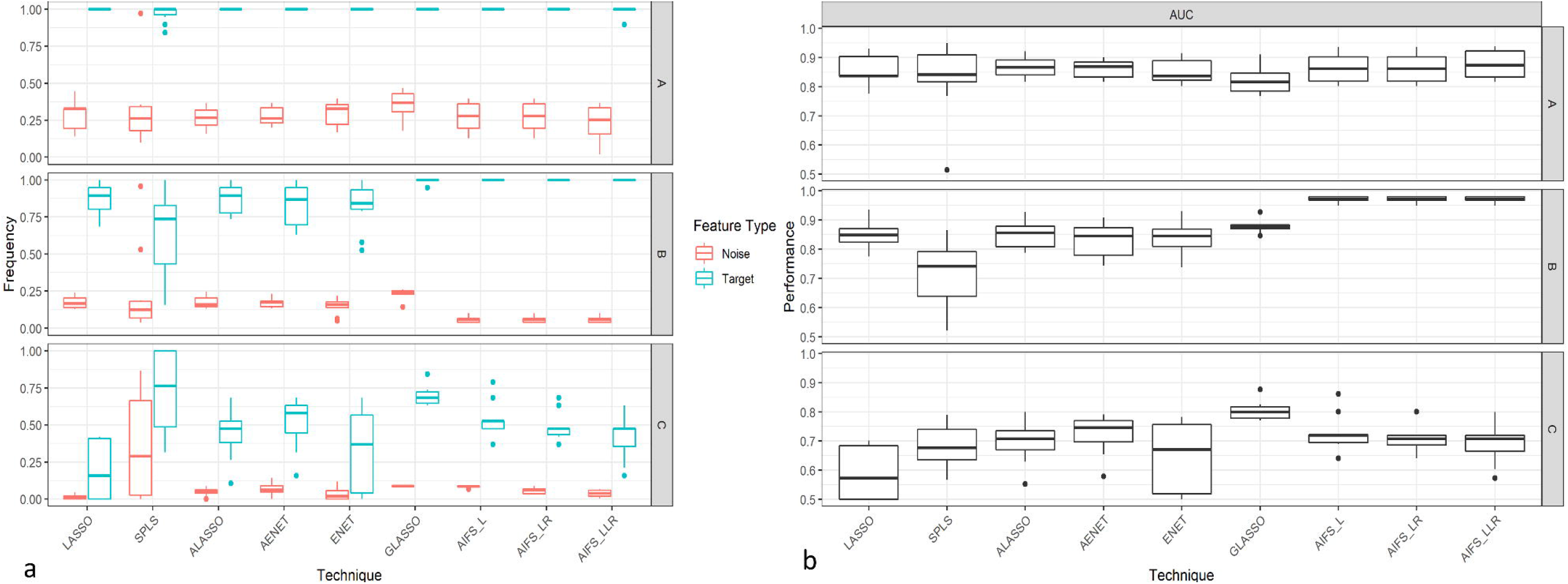
Feature selection performance comparison of different methods in interaction models a) Frequency of selection of target and noise features. b) AUC for predicting the target and noise features.

In AIFS, we used existing classic statistical techniques. The use of statistical techniques could have an important influence on the wrapper method performance [29]. However, a performance comparison between LASSO technique used in AIFS and as a standalone feature selection method clearly showed that AIFS could improve the LASSO performance. The AIFS performance suggests that the proposed methodology could enhance the feature selection performance of the existing statistical techniques by reducing the feature space and increasing the target feature percentage.

Table 5 shows the prediction performance of different methods. RMSE performance of the tested methods suggests that AIFS method performs consistently better or at par with the existing methods. In low dimensionality data (2_M, 4_M and 1_I), it is expected that all methods should give similar performance as standard methods are primarily developed for handling low dimensionality data, and results support it. AIFS method can provide better performance even in high dimensional settings (1_M and 3_M) and in the presence of interaction terms (2_I). However, at very high dimensional data (3_I), all methods perform poorly. These findings suggest that the AIFS may provide better or at par prediction performance than existing methods. Overall, the proposed method could expand the capability of existing techniques like non-penalized regression to operate in high-dimensional settings. However, computational intensiveness will be a significant limitation for the proposed methodology compared to standard methods. In summary, when we compare the performance of FS methods across different data dimensionality, performance of all methods deteriorates with an increase in data dimensionality, but performance of most standard methods decreases more drastically than AIFS.

**Table 5:** Outcome prediction performance of different approaches in simulated scenarios for the test dataset

| Methods | Performance (RMSE) |  |  |  |  |  |  |
| --- | --- | --- | --- | --- | --- | --- | --- |
|  | Marginal Model Scenarios |  |  |  | Interaction Model Scenarios |  |  |
|  | 1_M | 2_M | 3_M | 4_M | 1_I | 2_I | 3_I |
|  | Mean (95% Confidence Interval) |  |  |  |  |  |  |
| **ALASSO** | 0.44<br>(0.35-0.54) | 0.28<br>(0.23-0.33) | 0.39<br>(0.32-0.46) | 0.30<br>(0.26-0.35) | 0.44<br>(0.36-0.52) | 0.94<br>(0.74-1.13) | 1.36<br>(1.31-1.41) |
| **GLASSO** |  |  |  |  | 0.36<br>(0.3-0.43) | 0.65<br>(0.51-0.80) | **1.20**<br>**(1.15-1.26)** |
| **LASSO** | 0.45<br>(0.36-0.54) | 0.29<br>(0.24-0.34) | 0.40<br>(0.33-0.47) | 0.31<br>(0.26-0.36) | 0.40<br>(0.33-0.47) | 0.94<br>(0.76-1.13) | 1.36<br>(1.32-1.40) |
| **SPLS** | 0.45<br>(0.35-0.55) | 0.26<br>(0.21-0.31) | 0.43<br>(0.28-0.58) | 0.27<br>(0.23-0.31) | 0.52<br>(0.38-0.66) | 1.33<br>(1.21-1.45) | 1.47<br>(1.38-1.56) |
| **Enet** | 0.45<br>(0.36-0.53) | 0.29<br>(0.24-0.35) | 0.42<br>(0.34-0.5) | 0.32<br>(0.27-0.36) | 0.41<br>(0.34-0.49) | 1.02<br>(0.82-1.22) | 1.34<br>(1.29-1.38) |
| **AEnet** | 0.46<br>(0.35-0.57) | 0.28<br>(0.23-0.33) | 0.41<br>(0.33-0.48) | 0.31<br>(0.26-0.35) | 0.46<br>(0.38-0.54) | 0.97<br>(0.79-1.15) | 1.34<br>(1.30-1.39) |
| **AIFS-L** | 0.51<br>(0.38-0.65) | 0.28<br>(0.23-0.32) | 0.43<br>(0.34-0.52) | 0.31<br>(0.26-0.36) | **0.36**<br>**(0.29-0.43)** | **0.50**<br>**(0.40-0.61)** | 1.43<br>(1.30-1.57) |
| **AIFS-LLr** | **0.41**<br>**(0.26-0.56)** | **0.26**<br>**(0.21-0.31)** | **0.33**<br>**(0.27-0.39)** | **0.27**<br>**(0.22-0.32)** | 0.39<br>(0.31-0.48) | 0.58<br>(0.39-0.77) | 1.44<br>(1.33-1.55) |
| **AIFS-LR** | 0.46<br>(0.33-0.58) | 0.30<br>(0.26-0.33) | 0.34<br>(0.30-0.38) | 0.29<br>(0.26-0.33) | 0.56<br>(0.48-0.65) | 0.79<br>(0.68-0.91) | 1.35<br>(1.28-1.41) |

### Real Studies: Population Health Data

Four real studies are analyzed to evaluate the performance of AIFS and existing methods. Community Health Status Indicators (CHSI) study focuses on non-communicable diseases from US county with data (n=3141) containing 578 features [30] (Study I). National Social Life, Health and Aging Project (NSHAP) datasets focusing on the health and well-being of aged Americans contains multiple datasets. We chose two datasets (Study II and Study III) containing data for 4377 residents on 1470 features [31] and 3005 residents on 820 features [32]. Study IV is the Study of Women’s Health Across the Nation (SWAN), 2006-2008 dataset focusing on 887 *physical, biological, psychological and social* features in middle-aged women in the USA (n = 2245) [33].

The raw data of the real studies are processed for ease of analysis to obtain final cleaned datasets (Table 6). Features and samples are filtered to remove highly correlated features, non-continuous features, and missing values. Then, each dataset is randomly split into training and testing datasets. As the sample size is large, only 20% of data is used for training while remaining 80% of data is used for testing to create a high dimensional data setting. We compare the performance of different methods for marginal models and interaction models using mean RMSE of the test data in ten trials.

**Table 6:**
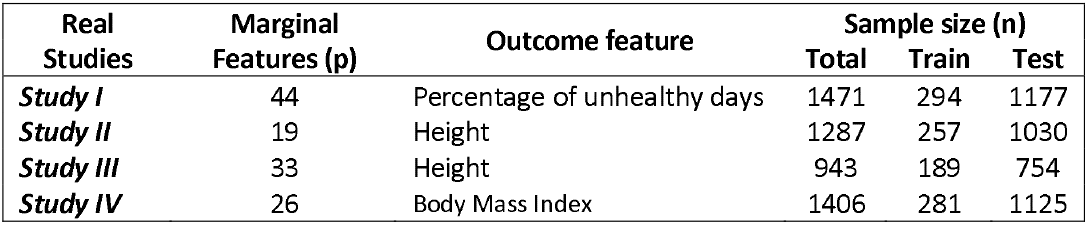
Summary of the real datasets

Table 7 summarizes the feature selection results. It is shown that standard methods are selecting a lesser number of features as compared to AIFS methods. However, the results from the previous simulated data studies suggest that standard methods may struggle to discriminate between target and noise features (Figure 1 and Figure 2). Further, the predictive performance results of AIFS method is better than the standard methods for both marginal as well as interaction models (Table 8). The better performance of the proposed method suggests that it may be more reliable than standard methods in identifying the target features.

**Table 7:**
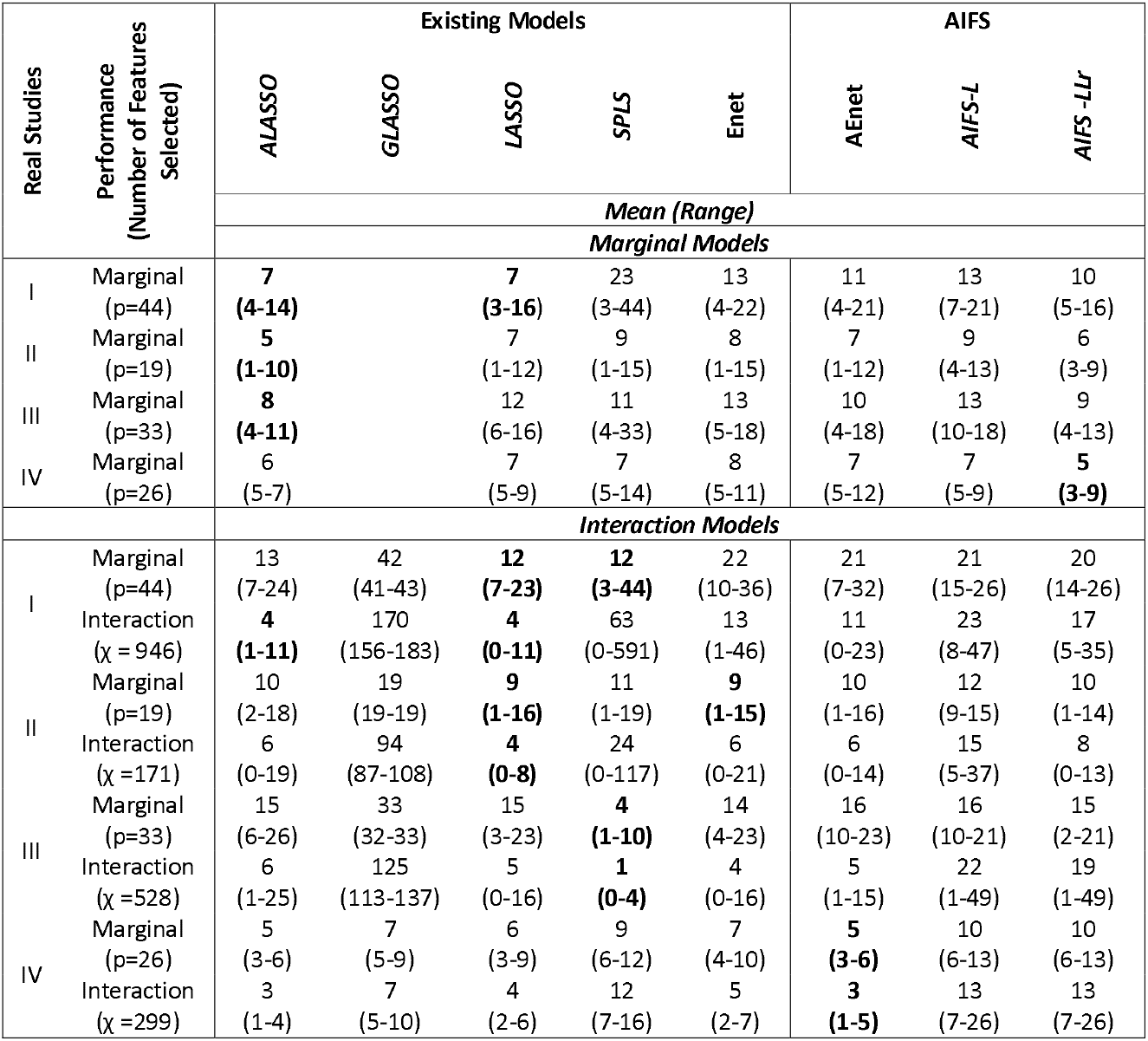
Number of features selected by different wrapper methods on the real studies

**Table 8:**
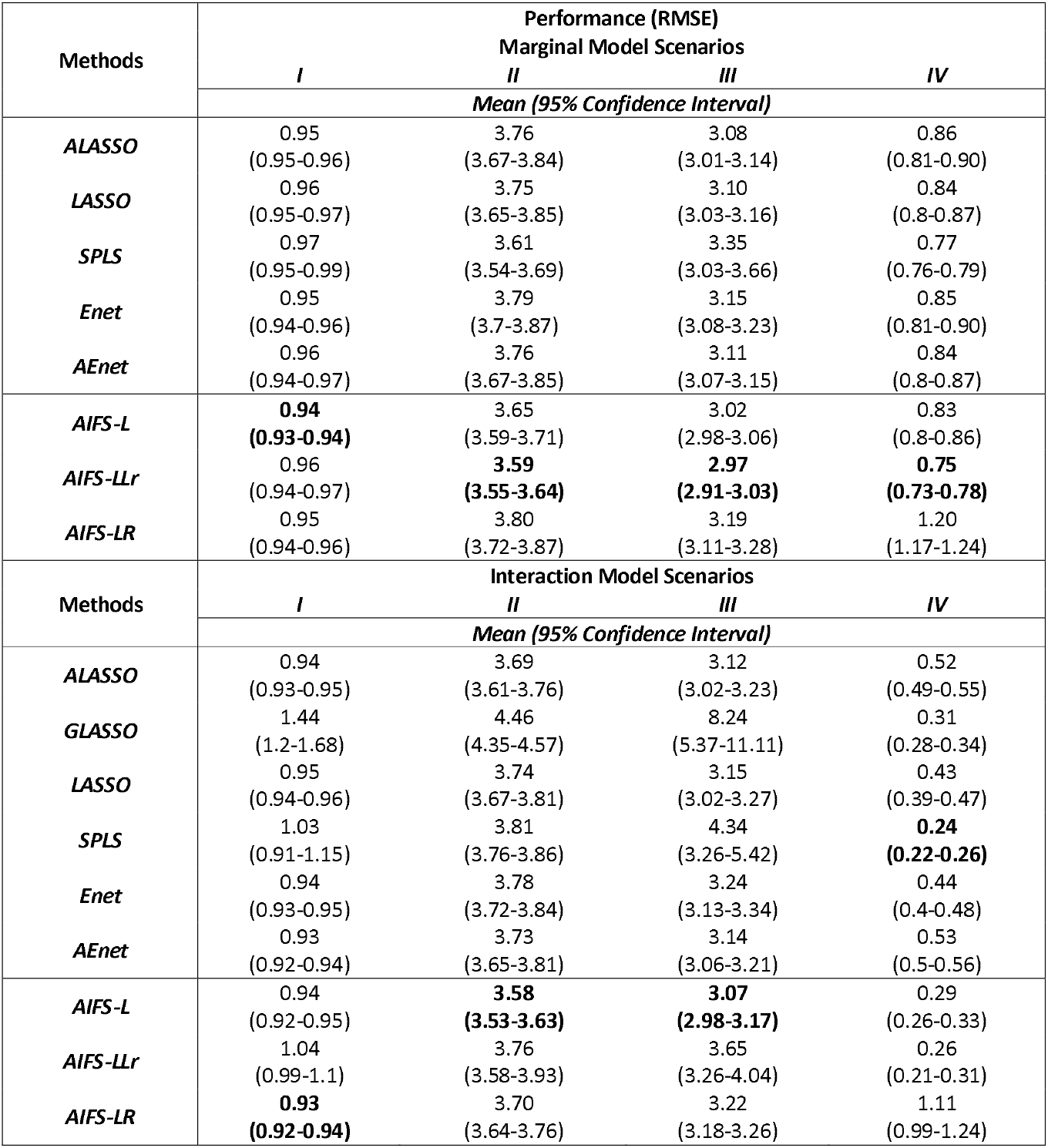
RMSE performance of different methods on the real studies for test data

| Methods | Performance (RMSE) |  |  |  |
| --- | --- | --- | --- | --- |
|  | Marginal Model Scenarios |  |  |  |
|  | *I* | *II* | *III* | *IV* |
|  | *Mean (95% Confidence Interval)* |  |  |  |
| **ALASSO** | 0.95<br>(0.95-0.96) | 3.76<br>(3.67-3.84) | 3.08<br>(3.01-3.14) | 0.86<br>(0.81-0.90) |
| **LASSO** | 0.96<br>(0.95-0.97) | 3.75<br>(3.65-3.85) | 3.10<br>(3.03-3.16) | 0.84<br>(0.8-0.87) |
| **SPLS** | 0.97<br>(0.95-0.99) | 3.61<br>(3.54-3.69) | 3.35<br>(3.03-3.66) | 0.77<br>(0.76-0.79) |
| **Enet** | 0.95<br>(0.94-0.96) | 3.79<br>(3.7-3.87) | 3.15<br>(3.08-3.23) | 0.85<br>(0.81-0.90) |
| **AEnet** | 0.96<br>(0.94-0.97) | 3.76<br>(3.67-3.85) | 3.11<br>(3.07-3.15) | 0.84<br>(0.8-0.87) |
| **AIFS-L** | **0.94**<br>**(0.93-0.94)** | 3.65<br>(3.59-3.71) | 3.02<br>(2.98-3.06) | 0.83<br>(0.8-0.86) |
| **AIFS-LLr** | 0.96<br>(0.94-0.97) | **3.59**<br>**(3.55-3.64)** | **2.97**<br>**(2.91-3.03)** | **0.75**<br>**(0.73-0.78)** |
| **AIFS-LR** | 0.95<br>(0.94-0.96) | 3.80<br>(3.72-3.87) | 3.19<br>(3.11-3.28) | 1.20<br>(1.17-1.24) |
| Methods | Interaction Model Scenarios |  |  |  |
|  | *I* | *II* | *III* | *IV* |
|  | *Mean (95% Confidence Interval)* |  |  |  |
| **ALASSO** | 0.94<br>(0.93-0.95) | 3.69<br>(3.61-3.76) | 3.12<br>(3.02-3.23) | 0.52<br>(0.49-0.55) |
| **GLASSO** | 1.44<br>(1.2-1.68) | 4.46<br>(4.35-4.57) | 8.24<br>(5.37-11.11) | 0.31<br>(0.28-0.34) |
| **LASSO** | 0.95<br>(0.94-0.96) | 3.74<br>(3.67-3.81) | 3.15<br>(3.02-3.27) | 0.43<br>(0.39-0.47) |
| **SPLS** | 1.03<br>(0.91-1.15) | 3.81<br>(3.76-3.86) | 4.34<br>(3.26-5.42) | **0.24**<br>**(0.22-0.26)** |
| **Enet** | 0.94<br>(0.93-0.95) | 3.78<br>(3.72-3.84) | 3.24<br>(3.13-3.34) | 0.44<br>(0.4-0.48) |
| **AEnet** | 0.93<br>(0.92-0.94) | 3.73<br>(3.65-3.81) | 3.14<br>(3.06-3.21) | 0.53<br>(0.5-0.56) |
| **AIFS-L** | 0.94<br>(0.92-0.95) | **3.58**<br>**(3.53-3.63)** | **3.07**<br>**(2.98-3.17)** | 0.29<br>(0.26-0.33) |
| **AIFS-LLr** | 1.04<br>(0.99-1.1) | 3.76<br>(3.58-3.93) | 3.65<br>(3.26-4.04) | 0.26<br>(0.21-0.31) |
| **AIFS-LR** | **0.93**<br>**(0.92-0.94)** | 3.70<br>(3.64-3.76) | 3.22<br>(3.18-3.26) | 1.11<br>(0.99-1.24) |

The results show that in Study III, marginal models performed better than their interaction models for all methods. Better performance of the marginal model compared to the interaction model suggests that AIFS cannot completely reject noise features and is sensitive to an increase in feature space. However, AIFS is still more robust than standard methods and can perform in different dimensions and datasets.

### Real Studies: Genomic Data

AIFS-L method is compared with StW method in the genomic datasets to determine the biological relevance of the solutions obtained from AIFS method. In many cancer studies, it is found that smoking can be detrimental to the cancer patient health [34, 35]. Further, an association between gene expression levels and cancer patient smoking habit has been reported [36]. Thus, it would be relevant to identify the genes in cancer patients which are associated with smoking-related traits. In this study, The Cancer Genomic Atlas (TCGA) program is used to get the data from nine cancer projects (Table 9) which maintained records related to amount smoked and gene expression profile of patients [37]. The sample size for these projects range from 89 to 592 samples with feature space *p* of 56602 genes. The gene expression profile is used as the input feature space and number of cigarettes smoked per day (CPD) is used as the outcome.

**Table 9:** Summary of the genomic datasets

| Datasets | Number of cigarettes smoked per day<br>($\mu(\sigma)$) | Sample Size<br>(n) | Feature Space<br>(p) |
| --- | --- | --- | --- |
| TCGA-BLCA | 1.16 (2.34) | 433 | 56602 |
| TCGA-CESC | 0.30 (0.62) | 307 | 56602 |
| TCGA-ESCA | 0.95 (1.21) | 172 | 56602 |
| TCGA-HNSC | 1.41 (1.89) | 544 | 56602 |
| TCGA-KICH | 0.21 (0.67) | 89 | 56602 |
| TCGA-KIRP | 0.42 (1.04) | 320 | 56602 |
| TCGA-LUAD | 1.53 (1.59) | 592 | 56602 |
| TCGA-LUSC | 2.44 (1.88) | 551 | 56602 |
| TCGA-PAAD | 0.46 (0.88) | 181 | 56602 |

Preliminary processing of all datasets is performed to reduce the input feature space and remove samples with missing values. The input feature space is reduced from 56602 to 50 features through multi-stage processing (Table 9). Step one involved removing the features which are not differentially expressed in cancer patients as compared to normal patients using *TCGAbiolinks* package [38]. Step two involved supervised dimensionality reduction of the differentially expressed genes using partial least squares technique and select top 100 features with highest absolute weights in first latent feature. Step three involved removing correlations among the features. Thus, among any pair of features with more than 0.8 absolute correlation, one feature is randomly selected. Step four involves selecting the top 50 features among the non-correlated features based on their absolute weight in the first latent feature obtained in step two. No interaction effects are considered for this analysis.

The performance of AIFS and StW in all datasets is compared on three metrics namely predictive performance, computation time and number of genes selected. The results are based on 10-fold cross-validation (Table 10). It observed that in all the datasets the predictive performance of AIFS based features is better or at par with StW based features. Further, it is observed that a smaller set of features are selected by AIFS as compared to StW which suggests AIFS could provide a more parsimonious set of features as compared to StW without compromising on the predictive performance of the features. In terms of computation time, the results are similar to those observed in simulation studies with StW taking less time than AIFS in most cases.

**Table 10:** Wrapper methods comparison of predictive performance, number of genes selected and computation time

| *Dataset* | Performance ($\mu$ [95% CI])<sup>*</sup> | | | | | |
| --- | --- | --- | --- | --- | --- | --- |
|  | Predictive Performance (RMSE) |  | Number of Genes Selected |  | Computation Time (minutes) |  |
|  | *StW* | *AIFS-L* | *StW* | *AIFS-L* | *StW* | *AIFS-L* |
| TCGA-BLCA | 0.79[0.31,1.27] | **0.78[0.30,1.26]** | 4[0,9] | **1[0,3]** | **5.9[3.2,8.6]** | 12.2[10.1,14.3] |
| TCGA-CESC | 1.00[0.84,1.16] | **0.98[0.84,1.13]** | 10[7,13] | **5[4,6]** | **11[7.7,14.2]** | 14.6[9.9,19.3] |
| TCGA-ESCA | 1.04[0.87,1.20] | **1.00[0.85,1.15]** | 11[5,17] | **8[2,14]** | **7.2[4.9,9.5]** | 27.9[3.6,52.2] |
| TCGA-HNSC | 0.99[0.82,1.16] | **0.98[0.81,1.15]** | 16[12,20] | **6[3,9]** | **11.4[8.7,14]** | 20.3[9.3,31.2] |
| TCGA-KICH | 1.03[0.61,1.46] | **0.82[0.39,1.25]** | 11[9,13] | **6[4,8]** | 50.2[24.7,75.7] | **10.6[7.5,13.7]** |
| TCGA-KIRP | 0.95[0.66,1.24] | **0.95[0.65,1.24]** | 19[18,20] | **15[11,19]** | **10.4[8.8,12]** | 41.1[12.5,69.8] |
| TCGA-LUAD | 1.02[0.93,1.11] | **1.02[0.94,1.09]** | 25[22,28] | **21[16,26]** | **11.6[9.1,14.1]** | 42.3[11.6,72.9] |
| TCGA-LUSC | 0.99[0.91,1.08] | **0.99[0.91,1.08]** | 2[1,3] | **1[0,2]** | **5.7[4.4,7]** | 12[8.8,15.2] |
| TCGA-PAAD | 1.26[0.74,1.79] | **1.24[0.75,1.73]** | 22[20,24] | **14[9,19]** | **10.8[7.6,14.1]** | 29[0.6,57.4] |

In order to assess the biological relevance of the genes selected by each method, selected genes of each dataset are pooled together to create final list of genes selected by each method. The results show that some genes are selected at a very high frequency in dataset during 10-fold feature selection process. Genes need to fulfill one of the two criteria of either having highest selection frequency or selection frequency of more than 80%. Accordingly, across nine datasets, AIFS provided 13 genes while StW provided 40 genes. 11 genes (VCX3A, WNT3A, CALHM5, ZMYND10, FOXE1, PLAT, BAAT, WFDC5, CGB5, FADD, APOE) are found to be common across the two methods. Among the 13 genes from AIFS method, seven genes (WNT3A [39], TMEM45A [40], BAAT [40], WFDC5 [41], HS3ST5 [42], CGB5 and APOE [43]) have been reported in literature to exert influence on tobacco or smoking-related traits. Further, AIFS identified six new genes (VCX3A, CALHM5, ZMYND10, FOXE1, PLAT, FADD) which could be related to smoking in cancer patients, thus providing an opportunity for identifying previously unknown biological functions.

## Discussion

Building models for each sample feature set obtained during the feature sampling stage of wrapper methods consume computational resources and may not always provide the best results. AIFS allows skipping the model building for many sample feature sets by training an AI model, i.e., the PPM model, which could predict the performance of sample feature sets. AIFS feature selection performance and predictive performance are better or at par than both the standard wrapper approach and penalized standard methods, namely LASSO, adaptive LASSO, group LASSO, Sparse PLS, Elastic net and adaptive elastic net.

The proposed method has certain limitations. The current study primarily focuses on testing the concept; thus, the study performed testing on limited datatypes. Future research could focus on evaluating the robustness of the approach using different types of data such as temporal data and categorical data, and outcomes such as binary outcomes and time to event outcomes. Other than data types, the focus could also be directed towards the algorithm used. Currently, the study uses a linear combination function for building actual models, but future studies could also explore the non-linear combination function for model building. Further, the current study reduced the need to build actual models in the wrapper approach but could not eliminate it. Therefore, future research could use other PPM building techniques like an artificial neural network and support vector machines to eliminate the need for actual models.

## Conclusion

In the paper, we propose AIFS, an innovative approach to perform wrapper based feature selection. The method is flexible enough to work with both marginal and interaction terms. The approach could be easily embedded with any of the wrapper techniques as it does not alter existing methods, which allows users to integrate the method in their existing wrapper pipelines. This approach could enhance the performance of existing wrapper techniques available in the literature for high dimensional datasets by accelerating the algorithm. AIFS can identify both the marginal features and interaction terms without using interaction terms in PPM, which could be critical in reducing the feature space an algorithm has to process.

The benefits of AIFS comes from using artificial intelligence to learn the dataset performance behavior and build the PPM, which replaces the actual model building process. The studies involving marginal effects with and without interaction effects in simulated data showed that AIFS could outperform existing methods in feature selection and prediction performance. Similar performance in real datasets also demonstrates the practical relevance of AIFS.

## Conceptual Framework

In a wrapper approach, given a dataset *D* of sample size *n* with *p* feature space and outcom *y*, a subset feature set *q* is created from *p*. In the standard wrapper approach (Figure 3a), a model is built for the subset of *D* containing *q* features and performance is estimated. This performance is used to select the next subset of *p*. This dependence of a standard wrapper approach upon model building step for each subset of feature to estimate its performance is targeted in our AIFS algorithm.

**Figure 3.**
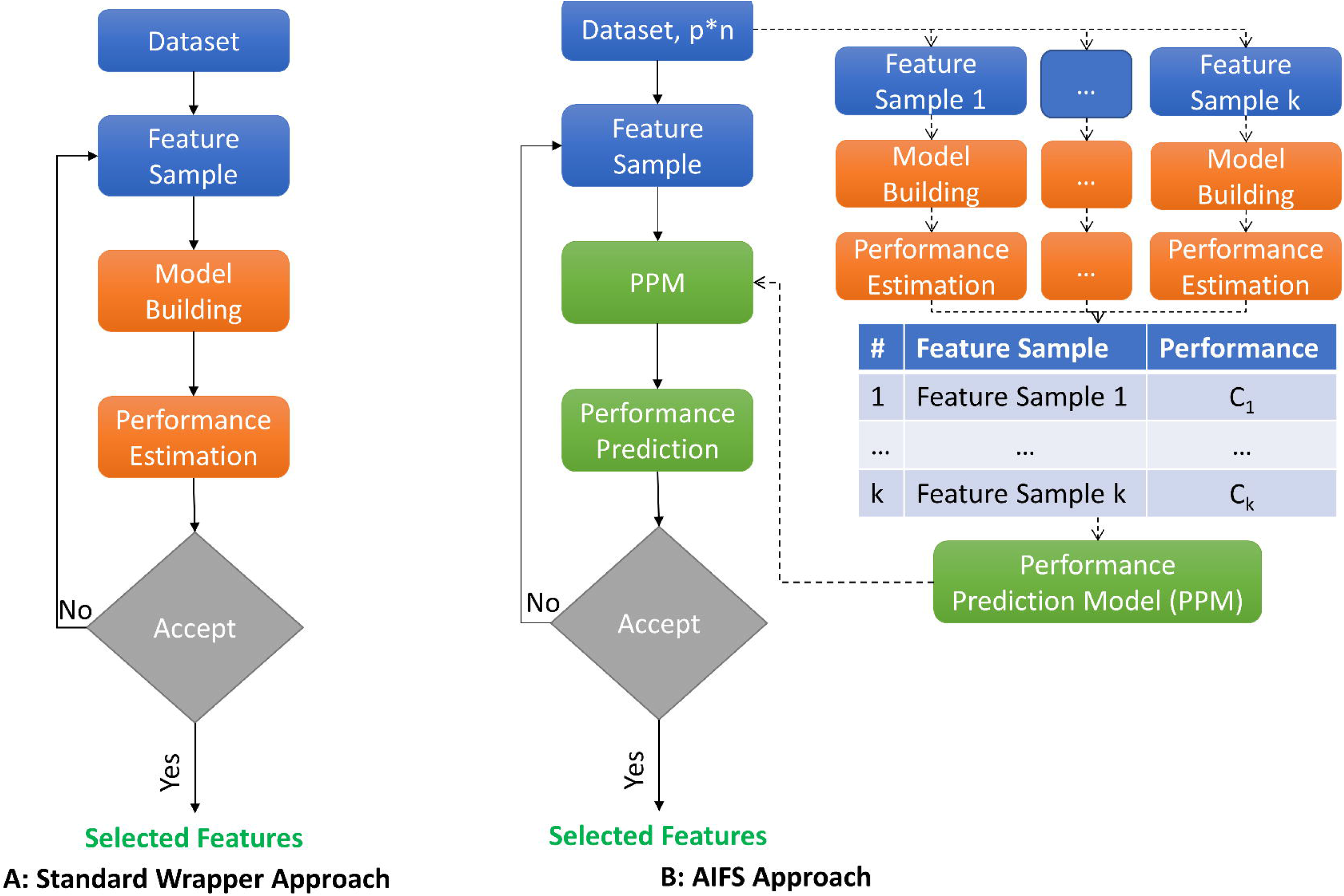
Flow chart of A) Standard wrapper approach and B) Proposed wrapper (AIFS) conceptual approach.

The conceptual framework used to design AIFS algorithm (Figure 3b) aims at reducing (or removing) the dependence of the wrapper algorithm on model building step for obtaining performance value of *q*. AIFS algorithm creates a random set 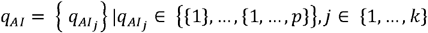 of Feature *k* samples, where each feature sample is a subset of *p*. The algorithm builds a model for *q*_*Ai*_ samples to estimate their performance *C* = {*C*_*j*_}. The algorithm creates a Performance Prediction Model (PPM) with *q*_*Ai*_ as the input and as the outcome using a machine learning model to enable performance prediction of any subset of *p*. Finally, the algorithm executes the standard wrapper approach, but uses PPM as a surrogate to the actual model building step that predicts rather than estimates the actual performance of *q*.

## Methodology

This section explains the design of AIFS algorithm based on the conceptual framework. The algorithm can be divided into four steps: performance prediction model, wrapper based coarse feature selection, embedded-feature selection and performance-based feature selection (Figure 4).

**Figure 4.**
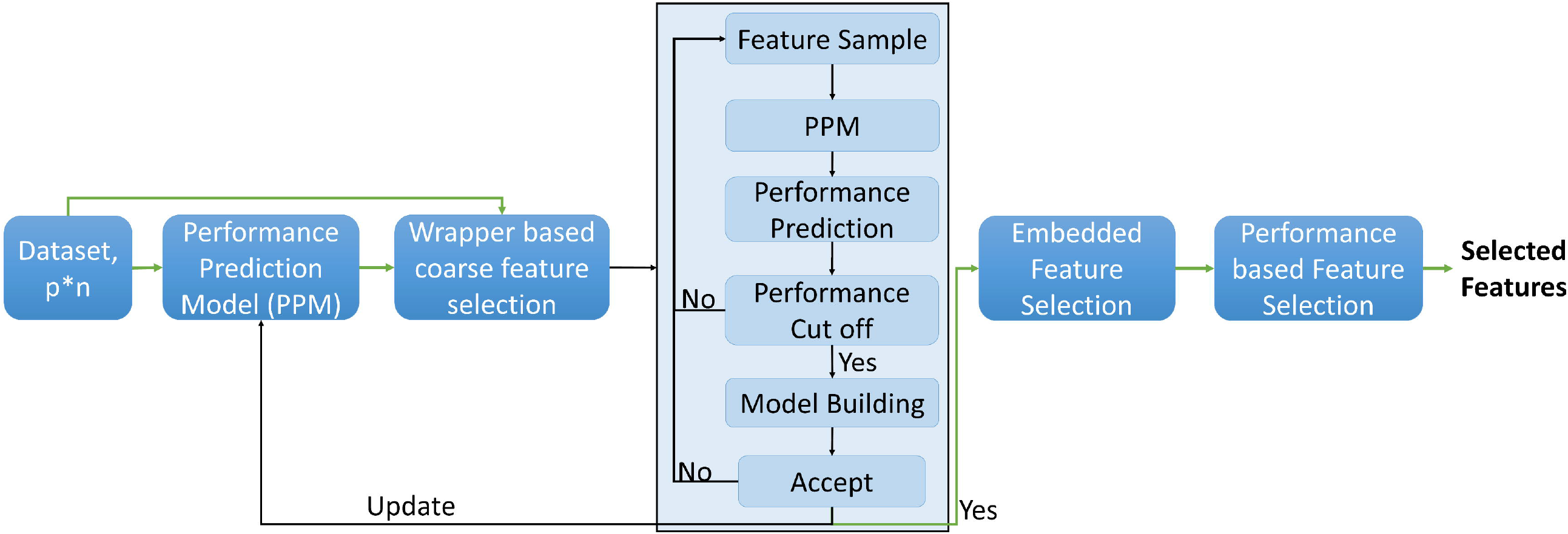
AIFS algorithm graphical flow chart. Dark Background represents main steps and light background represents sub-steps.

### Performance Prediction Model (PPM)

The algorithm generates random sample datasets containing 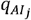 features, and sample size from *D*. A set of models *M* = {*m*_*j*_} are created from *k* sample datasets for an outcome, *y* using any modeling technique.

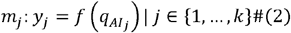

A performance set *C* = {*C*_*j*_} contains the performance of *M* models. The algorithm creates a performance dataset *D*_*pert*_, a matrix of features used in each model of *M* (*q*_*f*_) and their performance, *C*.

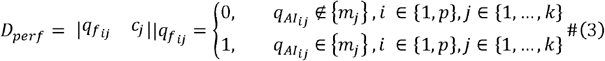

As shown in equation 3, feature matrix (*q*_f_) is a binary matrix that consists of *p* columns and rows. The matrix takes the value of 0 for *i*^*th*^column and *j*^*th*^ row, if *i*^*th*^ feature is not used in *m*_*j*_ model, else *i*^*th*^ column and row *j*^*th*^ takes the value of 1. PPM is constructed from *D*_*pref*_ to provide a predictive model for the outcome, using any machine learning technique.

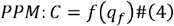

In this study, we have used LASSO to prepare *m*_*j*_ models and random forest to build the PPM. During the preliminary analysis (Additional File 1), it is found that predicted performance and actual performance is strongly and positively correlated, but predicted performance may not match the actual performance, as a result subset corresponding to best predicted performance may not be the best subset.

### Wrapper based coarse feature selection

The standard wrapper approach as shown in Figure 3a is an iterative process where a subset of feature is evaluated, and performance of the feature subset is used to select the next subset of features. In our work, we used genetic algorithm to search through the feature space iteratively as it is used in wide range of datasets [44–46]. In the proposed algorithm, we use PPM for all iterations to predict the performance *C*_*pred*_ of a feature set *q*. Since, we found that best *C*_*pred*_ may correspond to one of the high performing feature sets but not the best feature set, we validate *C*_*pred*_ values by building a model using *q* features to estimate the performance *C*_*true*_ (Figure 4). The algorithm uses user-defined criteria *val*_*crit*_ to select sample feature sets for validation of *C*_*pred*_ values.

In this study, the top quartile of C is used as the *val*_*crit*_ criterion, thus *q* with *C*_*pred*_ in top quartile of C are selected for model building. *D*_*pert*_ is updated with feature set *q* whose *C*_*true*_ value is available and consequently, is used to update PPM. The iteration stops when we get *q*_*wrap*_ features, which provide the best performance.

### Embedded feature selection

The *q*_*wrap*_ features obtained from the wrapper step are processed to obtain the final features because the prediction model does not explicitly provide the non-linear combinations of *q*_*wrap*_ features. Thus, an embedded feature selection model is used on *q*_*wrap*_ features for an outcome, *y* which allows the additional features χ like interactions terms to be incorporated. LASSO framework is used as the embedded model in the proposed algorithm.

### Performance-based feature selection

The features selected from the embedded model *q*_*embed*_ undergo the last stage of processing to provide final features *q*. This step selects features based on their contribution to the model

performance. *l* models 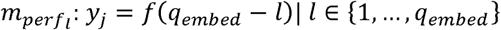 are prepared with each model containing *q*_*embed*_ −1 Features. *l* feature feature importance is determined from the 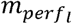 performance.

To obtain *l* feature robust importance, we create multiple models using bootstrapping of samples, and their performance 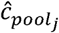 is pooled to get overall model performance 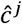. In this study, we use RIDGE regression for model building as we are focusing on high dimensional data and non-penalized linear regression could only work for cases with *q*_*embed*_ <*n*. Goodness of fit (R^2^) of out of the bag (OOB) samples is used as the performance metric. Finally, the performance metric is pooled to provide a coefficient of variation of R^2^ as the overall model performance for *l* feature.

A performance threshold *C*_*cutott*_ needs to be defined to select the features. Rather than using an arbitrary threshold, our algorithm uses a dynamic cutoff. The algorithm tries different performance feature space *q*_*best*_. In the current study, we use genetic algorithm to search through the thresholds and selects the threshold which provides the best performance *C*_*best*_ for the smallest performance threshold space. Two different techniques, namely non-penalized regression and adaptive RIDGE regression are used for the model building. Pseudo Algorithm summarizes the complete AIFS algorithm.

#### Pseudo Algorithm: AIFS

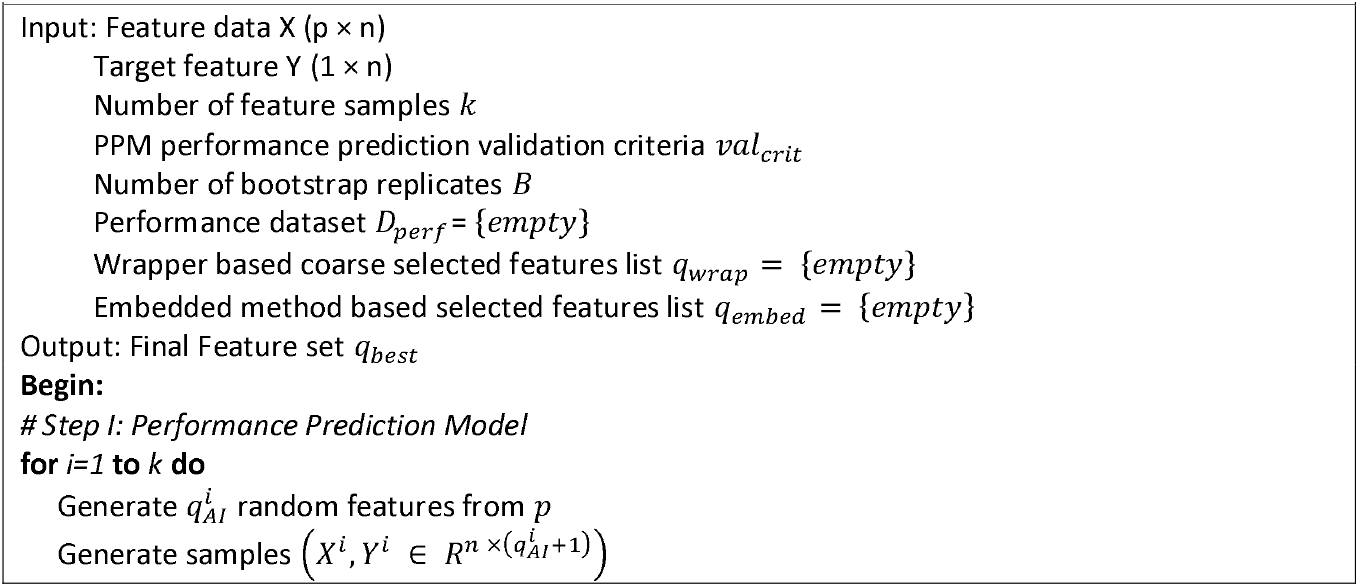
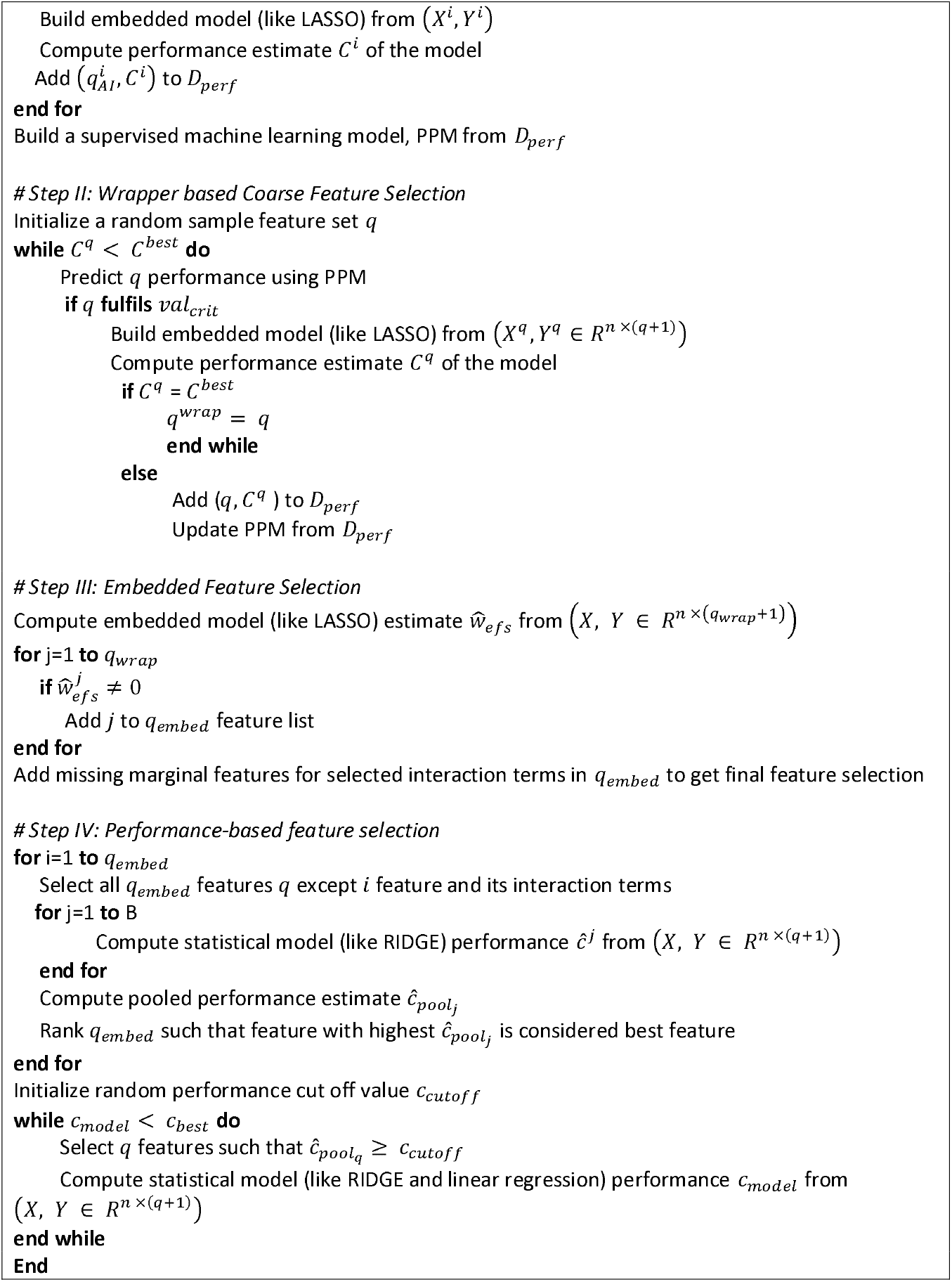

## List of abbreviations

AEnet: Adaptive Elastic Net
AI: Artificial Intelligence
AIFS: Artificial Intelligence infused wrapper based Feature Selection
ALASSO: Adaptive LASSO
AUC: Area Under Curve
CHSI: Community Health Status Indicators
Enet: Elastic Net
GLASSO: Group LASSO
NSHAP: National Social Life, Health and Aging Project
OOB: Out Of the Bag
PPM: Performance Prediction Model
RMSE: Root Mean Square Error
SPLS: Sparse Partial Least Squares
StW: Standard Wrapper
SWAN: Study of Women’s Health Across the Nation

## Declarations

### Ethics approval and consent to participate

Not Applicable

### Consent for publication

Not Applicable

### Availability of data and materials

All the datasets and code are in the github link: https://github.com/rahijaingithub/AIFS.

## Author Contributions

ALL AUTHORS HAVE READ AND APPROVED THE MANUSCRIPT.

**Conceptualisation:** RJ, WX

**Formal Analysis:** RJ

**Investigation:** RJ

**Methodology:** RJ, WX

**Software:** RJ

**Supervision:** RJ, WX

**Validation:** RJ, WX

**Writing-original draft:** RJ

**Writing-review & editing:** RJ, WX

## Acknowledgements

Not Applicable

