## Supplementary File 1 for "AIFS: A novel perspective, Artificial Intelligence infused wrapper based Feature Selection Algorithm on High Dimensional data analysis"

Appendix I:

Plot of actual performance vs predicted performance for the dataset with 50 features and 50 samples.

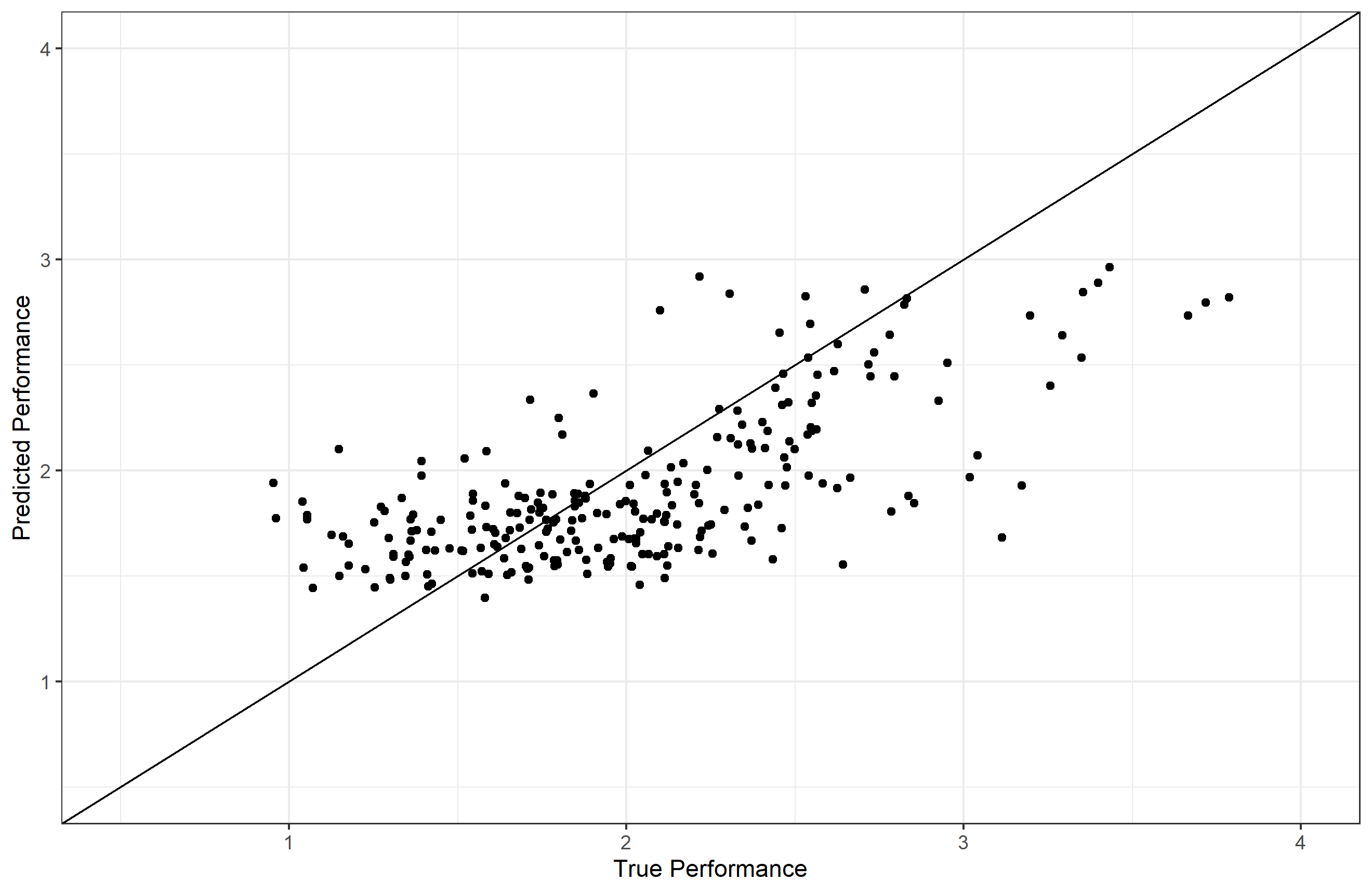

Appendix II:

Hyperparameters used in the AIWraFS algorithm are as follows:

| **#** | **Hyperparameter** | **Value** |
| --- | --- | --- |
| ***Performance Prediction Model (PPM)*** | | |
| 1 | k | 15n |
| ***Wrapper based coarse feature selection*** | | |
| 2 | Number of additional q added to dataset D to initiate PPM model retraining | 50 |
| ***Performance-based feature selection*** | | |
| 3 | Number of Bootstraps, B | 100 |
